# Map7D2 and Map7D1 facilitate microtubule stabilization through distinct mechanisms in neuronal cells

**DOI:** 10.1101/2021.10.27.466197

**Authors:** Koji Kikuchi, Yasuhisa Sakamoto, Akiyoshi Uezu, Hideyuki Yamamoto, Kei-ichiro Ishiguro, Kenji Shimamura, Taro Saito, Shin-ichi Hisanaga, Hiroyuki Nakanishi

## Abstract

Microtubule (MT) dynamics are modulated through the coordinated action of various MT-associated proteins (MAPs). However, the regulatory mechanisms underlying MT dynamics remain unclear. We show that the MAP7 family protein Map7D2 stabilizes MTs to control cell motility and neurite outgrowth. Map7D2 directly bound to MTs through its N-terminal half and stabilized MTs *in vitro*. Map7D2 localized prominently to the centrosome and partially on MTs in mouse N1-E115 neuronal cells, which expresses two of the four MAP7 family members, Map7D2 and Map7D1. Map7D2 loss decreased the resistance to the MT-destabilizing agent, nocodazole without affecting acetylated/detyrosinated stable MTs, suggesting that Map7D2 stabilizes MTs via direct binding. In addition, Map7D2 loss increased the rate of random cell migration and neurite outgrowth, presumably by disturbing the balance between MT stabilization and destabilization. Map7D1 exhibited similar subcellular localization and gene knock-down phenotypes to Map7D2. However, in contrast to Map7D2, Map7D1 was required for the maintenance of acetylated stable MTs. Taken together, our data suggest that Map7D2 and Map7D1 facilitate MT stabilization through distinct mechanisms in cell motility and neurite outgrowth.

## Introduction

Microtubule (MT) dynamics play crucial roles in a variety of cellular processes, including mitosis, and vesicle/organelle transport, as well as cell motility and morphology (Cleary & Hancock 2021, Etienne-Manneville 2013, Roll-Mecak 2020). MT dynamics are altered in response to various intrinsic or extrinsic signals and are then modulated through the coordinated actions of various MT-associated proteins (MAPs), which control the processes of dynamic instability (Cleary & Hancock 2021, Roll-Mecak 2020). Therefore, it is important to identify and characterize MAPs in order to understand the regulatory mechanisms of MT dynamics. We previously performed a comprehensive proteomic analysis of MT co-sedimented proteins from the brain and identified a series of functionally uncharacterized MT-binding proteins (Sakamoto et al 2008). The list included MAP7 family members Map7, Map7D1, and Map7D2, but not Map7D3. Among the MAP7 family, Map7 has been extensively characterized. Several lines of evidence suggest that Map7 has the ability to stabilize and reorganize MTs. Ectopic expression of Map7 induces MT bundling and resistance to nocodazole treatment-induced MT depolymerization (Masson & Kreis 1993). Map7 expression is upregulated during MT reorganization in response to the differentiation of keratinocytes (Fabre-Jonca et al 1999) and the establishment of apicobasal polarity in human colon adenocarcinoma cell lines, including Caco-2 and HT-29-D4 cells (Carles et al 1999, Masson & Kreis 1993). In addition, Map7 and the *Drosophila* Map7 homolog, Ensconsin (Ens), are involved in Kinesin-1-dependent transport by promoting the recruitment of a conventional Kinesin-1, Kif5b, and its *Drosophila* homolog, Khc, to MTs during various biological processes (Barlan et al 2013, Hooikaas et al 2019, Kikuchi et al 2018, Metzger et al 2012, Sung et al 2008, Tymanskyj et al 2018). The competition between Map7 and other MAPs for MT binding regulates the loading of motor proteins, thereby controlling the distribution and balance of motor activity in neurons (Monroy et al 2018, Monroy et al 2020). A considerable body of evidence has highlighted the important roles of Map7 in the regulation of MT dynamics. Similar to Map7, the MAP7 family member Map7D2 facilitates Kinesin-1-mediated transport in the axons of hippocampal neurons (Pan et al 2019), however, the function of Map7D2 in the regulation of MT dynamics and its relationship with other MAP7 family members remain unclear.

In this study, we first determined the tissue distribution and biochemical properties of Map7D2 in detail. Map7D2 is expressed predominantly in the glomerular layer of the olfactory bulb and the Sertoli cells of testes. Further, it directly associates with MTs through its N-terminal half, similarly to Map7, significantly enhancing MT stabilization. We also examined the cellular functions of Map7D2 using the N1-E115 mouse neuroblastoma cell line, which expresses both Map7D2 and Map7D1, but not Map7 nor Map7D3. Map7D2 predominantly localizes to the centrosome and partially on MTs, and suppresses cell motility and neurite outgrowth by facilitating MT stabilization via direct binding. Finally, we determined the functional differences between Map7D2 and Map7D1 with regard to MT stabilization in N1-E115 cells. Although Map7D1 exhibits similar subcellular localization and gene knock-down phenotypes to Map7D2, Map7D1 is required to maintain the amount of acetylated tubulin in contrast to Map7D2. These results suggest that Map7D2 and Map7D1 facilitate MT stabilization through distinct mechanisms for the control of cell motility and neurite outgrowth.

## Results

### Map7D2 is highly expressed in the glomerular layer of the olfactory bulb and the Sertoli cells of testes

In our previous comprehensive proteomic analysis of co-sedimented MT proteins from the rat brain, we identified a number of novel proteins (Sakamoto et al 2008). In the present study, we focused on Map7D2, which is one of the MAP7 family members (Fig. S1A). Although the tissue distribution of Map7 has been well-analyzed by using mice (Fabre-Jonca et al 1998, Komada et al 2000), that of Map7D2 has not been analyzed. To analyze the tissue distribution of Map7D2, we first performed northern blotting analysis using total RNA extracted from various rat tissues. Northern blotting analysis showed that the approximately 4.2-kb mRNA was hybridized only in the brain and testis, being more abundant in the former (Fig. 1A). Of note, no detectable signal was observed in other rat tissues examined, including the heart, spleen, lung, liver, skeletal muscle, and kidney.

**Figure 1.**
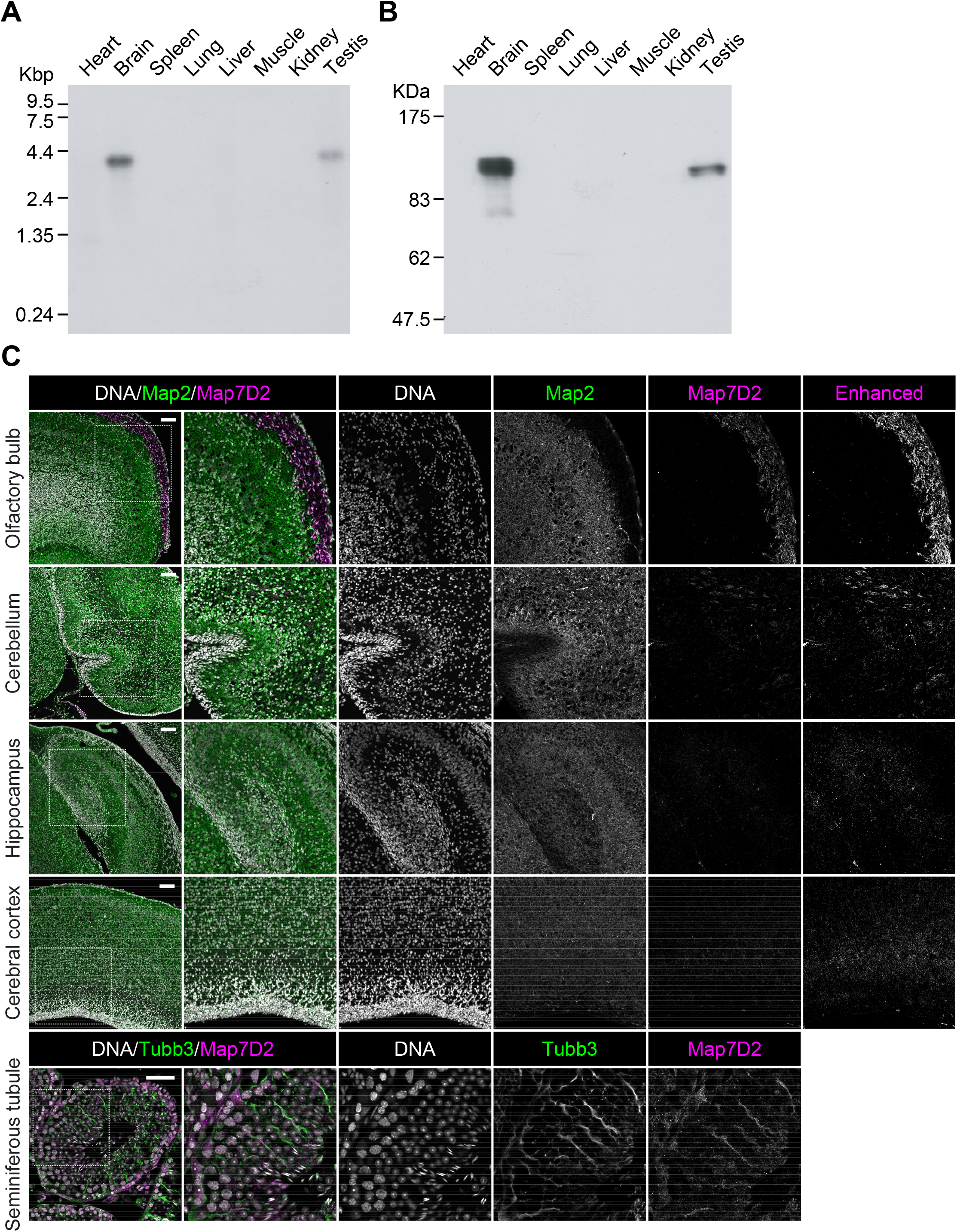
Tissue distribution of Map7D2. **(A)** The full scan image of Northern blot analysis. An RNA blot membrane (CLONTECH) was hybridized with the ^32^P-labeled full coding sequences of rMap7D2 according to the manufacturer’s protocol. **(B)** The full scan image of Immunoblotting analysis. Various tissue lysates (20 μg of protein) were subjected to SDS-PAGE, followed by immunoblotting with the anti-Map7D2 antibody. **(C)** Expression patterns of Map7D2 in the brain and testis by immunofluorescence. Upper panels, frozen sagittal sections of postnatal day 0 mouse brains were stained with anti-Map7D2 (magenta) and antibodies against mature neuron marker Map2 (green). DNA was labeled with DAPI (gray). For a comparison of signal intensities, images were captured under the same parameters. Contrast-enhanced images of Map7D2 staining were shown in the rightmost column. Lower panels, frozen coronal sections of adult mouse testis were stained with anti-Map7D2 (magenta) and antibodies against Sertoli cell marker Tubb3 (green). DNA was labeled with DAPI (gray). Data information: In **(C)**, scale bars in upper panels or lower panels are represented as 100 or 50 μm, respectively.

Next, we investigated the tissue distribution of Map7D2 at the protein level by immunoblotting. For the immunoblotting analysis, we raised an anti-Map7D2 polyclonal antibody using aa 1-235 of rat Map7D2 (rMap7D2) as an epitope (Fig. S2A). Using lysates from HeLa cells transfected with an empty vector, hMap7-V5His_6_, or rMap7D2-V5His_6_, we confirmed that the antibody detected Map7D2, but not Map7 (Fig. S2A). In addition, we evaluated antibody specificity by siRNA-mediated knock-down of endogenous *Map7d2*. For this experiment, we used a mouse neuroblastoma cell line, N1-E115, in which the expression of *Map7d2* and *Map7d1*, but not *Map7* and *Map7d3*, was detected by quantitative real-time PCR (RT-qPCR) (Fig. S2B). We designed three independent siRNAs against *Map7d2* or *Map7d1*. The immunoreactive band disappeared following treatment with each *Map7d2* siRNA, but not the control or *Map7d1* siRNA (Fig. S2C), indicating that the antibody specifically recognized Map7D2. We then performed immunoblotting analysis using lysates from various rat tissues. Consistent with the northern blotting analysis, Map7D2 was detected at the protein level only in the brain and testis, while no immunoreactive bands were detected in other rat tissues (Fig. 1B). In contrast to Map7D2, Map7 is widely expressed in a variety of mouse organs that contain epithelium (Fabre-Jonca et al 1998, Komada et al 2000). These data suggest that the mechanism of expression regulation may vary between Map7D2 and Map7.

We further analyzed the expression patterns of Map7D2 in the brain and testis by immunofluorescence. Based on RNA-seq CAGE, RNA-Seq, and SILAC database analysis (Expression Atlas, https://www.ebi.ac.uk/gxa/home/), Map7D2 expression was detected in the cerebellum, hippocampus, and olfactory bulb, and not in the cerebral cortex (Fig. S3). We confirmed Map7D2 expression in the above four brain tissue regions of postnatal day 0 mice by immunofluorescence. Among these regions, Map7D2 was the most highly expressed in the Map2-negative area of the olfactory bulb, *i*.*e*., the glomerular layer (Fig. 1C). Weak signals were detected in the cerebellum, and marginal signals were observed in the hippocampus and cerebral cortex (Fig. 1C). In contrast to Map7D2, Map7 has not been reported to be expressed in the glomerular layer of the olfactory bulb, while in the nervous system, Map7 is strongly expressed in the ganglia (Fabre-Jonca et al 1998, Komada et al 2000). Next, we analyzed Map7D2 expression in the seminiferous tubules of adult mice. Map7D2 signals were merged with Tubb3 signals, a marker for Sertoli cells (Fig. 1C), indicating that similar to Map7 (Komada et al 2000), Map7D2 is expressed predominantly in Sertoli cells. Taken together, these data suggest that *in vivo*, Map7D2 may function in the glomerular layer of the olfactory bulb and the Sertoli cells of the testis.

### Map7D2 has an ability to stabilize MTs

MAP7 family members share two conserved regions (Fig. S1A-C). The amino acid sequences of the N-terminal (aa 53-138) and C-terminal (aa 389-562) regions of human Map7D2 (hMap7D2) were 64.0% and 42.9% identical to those of human Map7 (hMap7), respectively, while other regions showed no significant homology to hMap7 (Fig. 2A). Using the rat brain cDNA library, we obtained rMap7D2 cDNA by PCR. The cloned cDNA encoded a protein consisting of 763 aa with a molecular weight of 84,823 (DDBJ/EMBL/GenBank accession number AB266744) (Fig. 2A). The full-length aa sequence of rMap7D2 was 68.1% identical to that of hMap7D2. For subsequent experiments, we used the rMap7D2 that we cloned.

**Figure 2.**
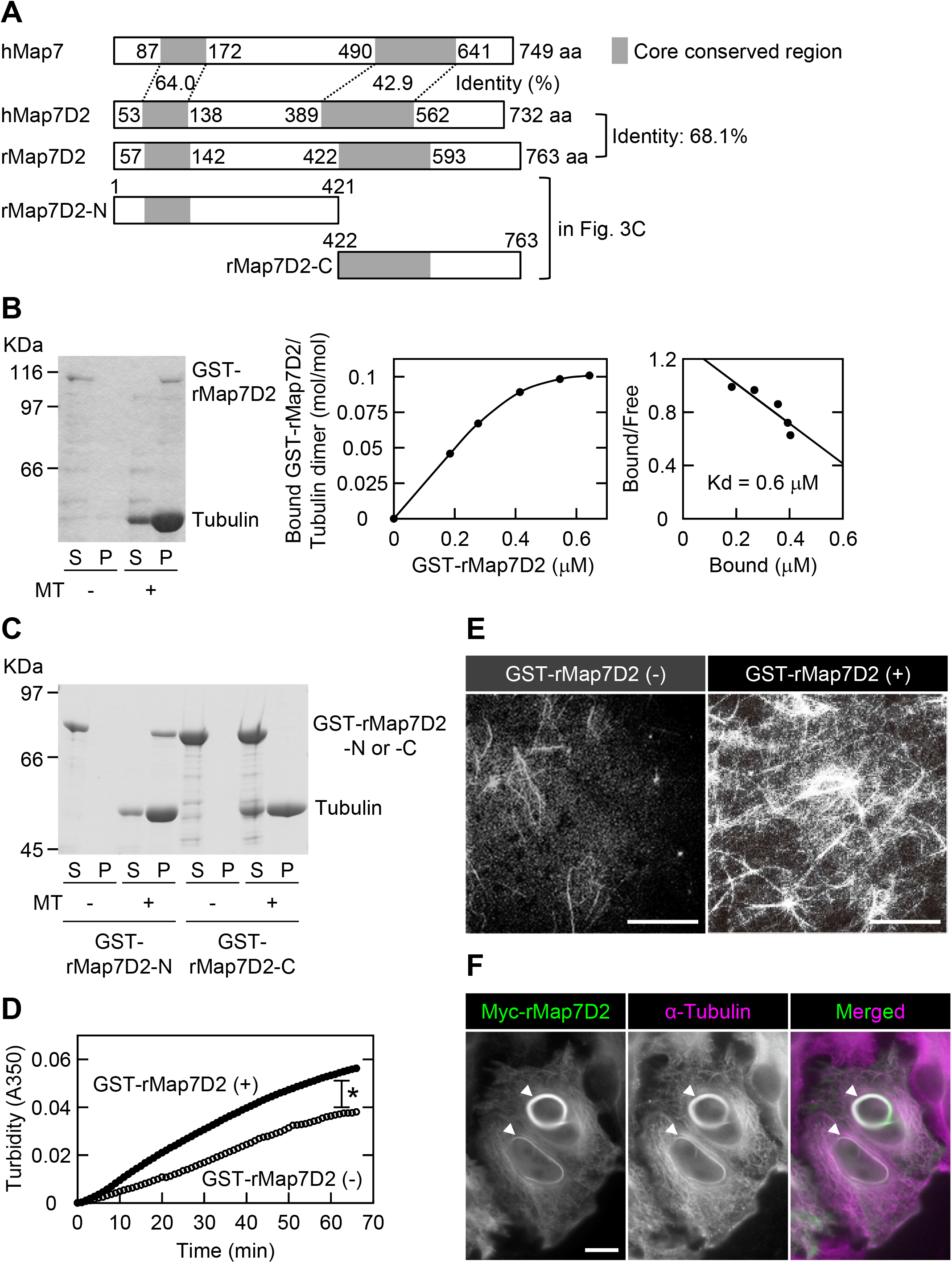
Map7D2 has the ability to stabilize MTs. (**A**) Schematic structures of hMap7, hMap7D2, and rMap7D2. (**B**) Co-sedimentation of rMap7D2 with MTs. Left panel, GST-rMap7D2 (34 μg/mL) was mixed with MTs, followed by ultracentrifugation. Comparable amounts of the supernatant and pellet fractions were subjected to SDS-PAGE, followed by CBB protein staining. S, supernatant; P, pellet. Middle panel, various amounts of GST-rMap7D2 were mixed with MTs, followed by ultracentrifugation. Amounts of free and bound GST-rMap7D2 were calculated by determining protein amounts from the supernatant and pellet fractions, respectively, with a densitometer. Right panel, Scatchard analysis. (**C**) Location of the MT-binding domain. GST-rMap7D2-N (80 μg/mL) or GST-rMap7D2-C (200 μg/mL) was mixed with MTs, followed by ultracentrifugation. Comparable amounts of the supernatant and pellet fractions were subjected to SDS-PAGE, followed by CBB protein staining. S, supernatant; and P, pellet. (**D**) Turbidity measurement. GST-rMap7D2 was mixed with tubulin. The sample was incubated at 37 □ and continuously monitored at 350 nm using a spectrophotometer. (○) without GST-rMap7D2; and (●) with GST-rMap7D2. (**E**) Immunofluorescent observation. GST-rMap7D2 was incubated for 20 min at 37 □ with rhodamine-labeled tubulin. After fixation, the sample was spotted on a slide glass and viewed under a fluorescence microscope. (**F**) HeLa cells transiently overexpressing Myc-rMap7D2. Myc-rMap7D2 was transfected into HeLa cells, and the cells were then double-stained with anti-Myc and anti-α-tubulin antibodies. Arrowheads show MT bundles. Data information: In **(D)**, *, *P* < 0.003 (the F-test). Scale bars, 50 μm in **(E)** and 10 μm in **(F)**.

We sought to determine whether rMap7D2 directly binds to MTs. To this end, we performed an MT co-sedimentation assay using recombinant rMap7D2. When GST-rMap7D2 was incubated with MTs, followed by ultracentrifugation, it was recovered with MTs in the pellet (Fig. 2B). The dissociation constant (Kd) was calculated to be approximately 6 × 10^−7^ M (Fig. 2B). This value is comparable to those of the well-known MAPs Tau and CLIP-170 (Gustke et al 1994, Lansbergen et al 2004). The stoichiometry of GST-rMap7D2 binding to tubulin was calculated to be one GST-rMap7D2 molecule per about ten tubulin α/β heterodimers. This value was also comparable to that of Map7 (Bulinski & Bossler 1994). It has been reported that Map7 binds to MTs through a conserved region on the N-terminal side, while Map7D3 binds via a conserved region on not only the N-terminal but also the C-terminal sides (Sun et al 2011, Yadav et al 2014). To further examine the location of the MT-binding domain of rMap7D2, the N-terminal (aa 1-421) and C-terminal (aa 422-763) halves were subjected to an MT co-sedimentation assay (Fig. 2A). The N-terminal half was co-sedimented with MTs, whereas the C-terminal half was not (Fig. 2C). These results indicate that the MT-binding domain of rMap7D2 is located only at the N-terminal half, similarly to that of Map7, but not Map7D3. Meanwhile, a region within the C-terminus of Map7 is required for complex formation with Kif5b, the heavy chain of Kinesin-1 (Fig. S4), and is involved in Kif5b-dependent transport by loading Kif5b onto MTs (Hooikaas et al 2019, Kikuchi et al 2018, Metzger et al 2012, Tymanskyj et al 2018). Map7D2 also formed a complex with Kif5b (Fig. S4), as previously reported (Pan et al 2019). Taken together, these data suggest that the biochemical properties are largely conserved between Map7D2 and Map7.

Next, we tested whether rMap7D2 affects MT assembly. The MT turbidity assay was used to analyze the effect of rMap7D2 on the kinetics of MT assembly. The addition of rMap7D2 significantly enhanced the amount of polymerized MTs in a time-dependent manner, whereas tubulin self-polymerized even in the absence of rMap7D2 (Fig. 2D). Identical results were observed by fluorescence microscopy analysis using rhodamine-labeled tubulin (Fig. 2E). Furthermore, we investigated the ability of Map7D2 to bundle MTs in HeLa cells. Consistent with the *in vitro* data, overexpression of Myc-rMap7D2 induced MT bundling in HeLa cells (Fig. 2F). Taken together, these results indicate that Map7D2 facilitates MT stabilization.

### Map7D2 localizes prominently to the centrosome and partially to MTs

Following the biochemical characterization of Map7D2, we sought to determine its functions within the cell. To this end, we used N1-E115 cells that express Map7D2 and Map7D1 (Fig. S2B and C). First, we analyzed the subcellular localization of Map7D2 in N1-E115 cells. N1-E115 cells can undergo neuronal differentiation in response to DMSO under conditions of serum starvation (Kimhi et al 1976), and most of the cells extend neurites up to 12 h after treatment with 1% DMSO (Fig. S5A) (Smit et al 2003). In both proliferating and differentiated cells, Map7D2 localized prominently to the centrosome and partially to MTs (Fig. 3A-C). These localizations were confirmed in N1-E115 cells stably expressing EGFP-rMap7D2 (Fig. 3D and E). Furthermore, during cytokinesis, Map7D2 accumulated at the midbody, where MT bundling occurs (Fig. 3B). Similarly, localization of Map7D2 was also observed at neurites, where MT bundling is also known to occur (Fig. 3C). Together with the biochemical properties, these subcellular localization data suggest that Map7D2 is involved in MT stabilization within the cell.

**Figure 3.**
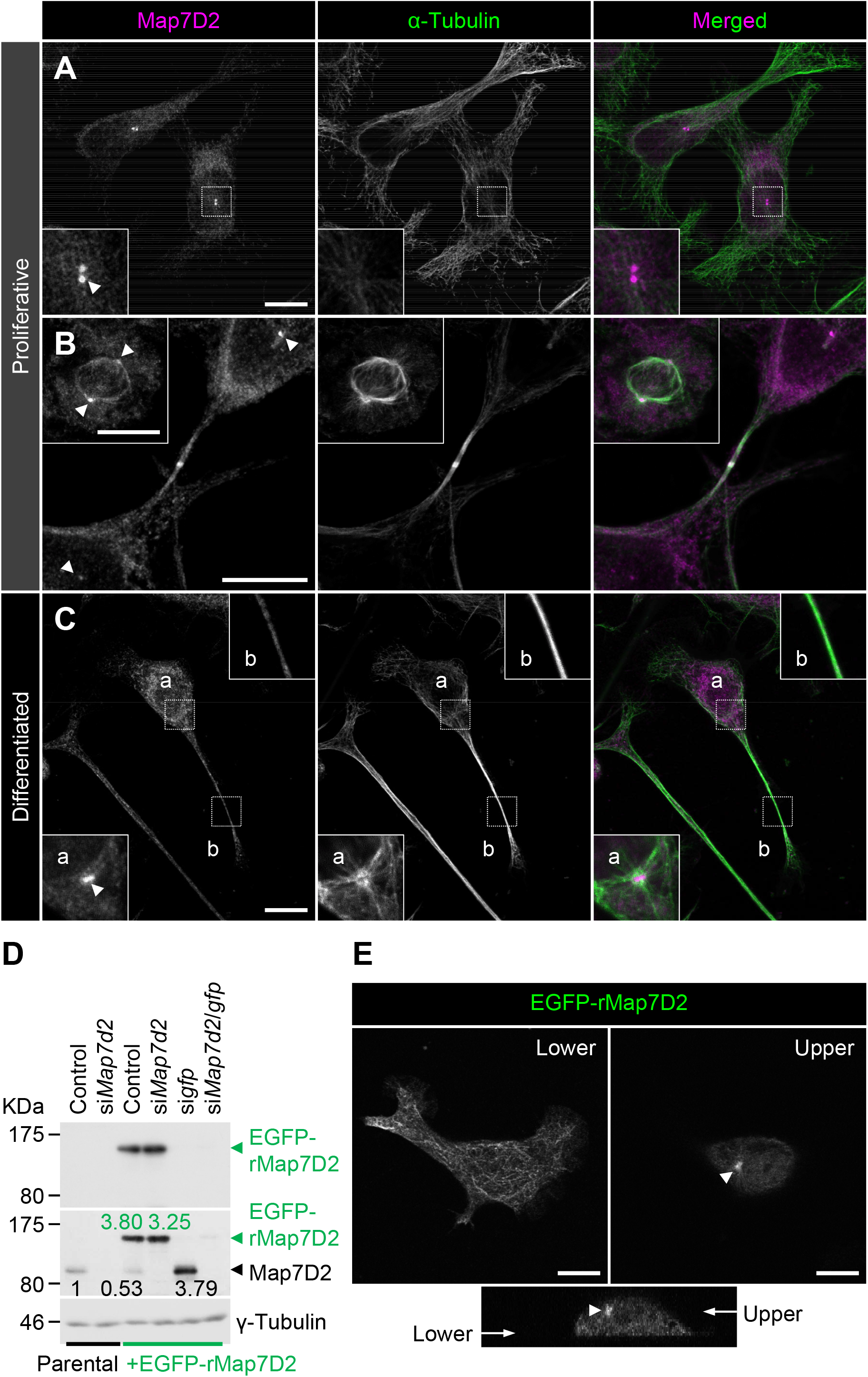
Subcellular localization of Map7D2 in proliferative and differentiated N1-E115 cells. (**A-C**) Localization of Map7D2 in interphase (**A**), mitosis (**B**), and differentiation state (**C**) of N1-E115 cells. Cells were double-stained with anti-Map7D2 and anti-α-tubulin antibodies. In **A**, the insets show enlarged images of regions indicated by a white box. In **B**, the inset shows metaphase cells. In C, images of differentiated cells were captured by z-sectioning, because the focal planes of the centrosome and neurites are different. Each inset shows an enlarged image of the region indicated with a white box at each focal plane. Arrowheads indicate the centrosomal localization of Map7D2. (**D**) Generation of N1-E115 cells stably expressing EGFP-rMap7D2. To check the expression level of EGFP-rMap7D2, lysates derived from the indicated cells were probed with anti-GFP (top panel) and anti-Map7D2 (middle panel) antibodies. The blot was reprobed for γ-tubulin as a loading control (bottom panel). The amount of endogenous Map7D2 or EGFP-rMap7D2 was normalized to the amount of γ-tubulin, and the value relative to endogenous Map7D2 in the parental control was calculated. (**E**) Confirmation for subcellular localization of Map7D2 using N1-E115 cells stably expressing EGFP-rMap7D2. Images were captured by z-sectioning. Top panels show images taken with the lower or upper focal plane, and bottom panels show the image reconstructed in the z-axis direction. Arrow head shows centrosomal localization of Map7D2. Data information: Scale bars, 10 μm in **(A-C, E)**.

As N1-E115 cells express another Map7 family member, Map7D1, we also determined its subcellular localization. Map7D1 exhibited similar localization to that of Map7D2 in both proliferative and differentiated states (Fig. S5B-D). In the previous analysis using HeLa cells (Hooikaas et al 2019, Kikuchi et al 2018), Map7D1 localizes to MTs, especially from the vicinity of the centrosome toward the MT plus-end, with gradual weakening. This pattern is slightly different from the localization pattern in N1-E115 cells, suggesting that the mechanisms regulating localization patterns of Map7D2 and Map7D1 may differ depending on the cell type. Interestingly, *Map7d1* knock-down upregulated Map7D2 expression, as confirmed with three different siRNAs (Fig. S2C), suggesting that Map7D2 expression is affected by changes in Map7D1 expression, not by off-target effects of a particular siRNA. Moreover, the amount of endogenous Map7D2 was decreased to 53% in N1-E115 cells stably expressing EGFP-rMap7D2 (Fig. 3D), suggesting that EGFP-rMap7D2 expression suppresses endogenous Map7D2 expression. In this cell line, the total amount of Map7D2 was increased, however, when EGFP-rMap7D2 was depleted using si*gfp* in this cell line, endogenous Map7D2 was expressed to the same level as EGFP-rMap7D2 before knock-down (Fig. 3D). Together with the finding that *Map7d1* knock-down increased the amount of Map7D2, these findings indicate that the amount of Map7D2 in the cells is regulated in response to the amount of Map7D1 and exogenous Map7D2. In contrast, *Map7d2* knock-down did not affect Map7D1 expression (Fig. S2C), and identical results were observed in the *Map7d2* knock-out (*Map7d2*^*-/-*^) N1-E115 cells we generated (Fig. S6A and B). As Ma7D2 expression was upregulated upon suppression of Map7D1 expression, Map7D2 has the potential to functionally compensate for Map7D1 loss. Therefore, we decided to analyze the phenotypes of single and double knock-downs for *Map7d2* and *Map7d1* in the following experiments.

### Map7D2 and Map7D1 facilitate MT stabilization through distinct mechanisms for the control of cell motility and neurite outgrowth

Our biochemical analyses led to the possibility that Map7D2 is involved in MT stabilization within the cell. We initially analyzed the effects of *Map7d2* or *Map7d1* knock-down on the resistance to nocodazole, a MT-destabilizing agent. In control N1-E115 cells, a small degree of MT shrinkage was observed, and elongated MT arrays were retained even after treatment with a low concentration of nocodazole for 1 h (Fig 4A). In contrast, not only *Map7d2* but also *Map7d1* knock-down dramatically increased MT shrinkage when subjected to nocodazole treatment at the same concentration (Fig 4A), indicating that both Map7D2 and Map7D1 are required for MT stabilization within the cell. Further, to investigate the possibility that Map7D2 or Map7D1 is involved in MT elongation, we measured the amount of EB1-decorated MTs at the cell periphery based on the intensity of EB1. The knock-down of either *Map7d2* or *Map7d1* did not affect the intensity of EB1 at the cell periphery, compared to control N1-E115 cells (Fig 4B), suggesting that both Map7D2 and Map7D1 are dispensable for the elongation of EB1-decorated MTs. Since Map7D2 and Map7D1 can form a complex with Kif5b (Fig. S4) (Kikuchi et al 2018, Pan et al 2019), we also examined whether *Map7d2* or *Map7d1* knock-down affects the distribution of Kif5b foci. Even after *Map7d2* or *Map7d1* knock-down, the distribution of Kif5b foci was similar to that in control N1-E115 cells (Fig. 4B). Kif5b foci were predominantly located at the internal regions of the cell (Fig. 4B), and some were partly observed in the protrusions (Fig. S7A). Taken together, these data indicate that Map7D2 and Map7D1 primarily stabilize MTs in N1-E115 cells. Considering the possibility that the Map7D2 dynamics are altered when MT stability is changed, e.g., before and after differentiation induction, we analyzed the Map7D2 dynamics at the centrosome by fluorescence recovery after photobleaching (FRAP) using N1-E115 cells stably expressing EGFP-rMap7D2. Compared to the proliferative state, the recovery rate of EGFP-rMap7D2 was reduced, and the immobile fraction of Map7D2 was increased in the differentiated state (Fig. 4C and Fig. S7B). These data suggest that an increase in immobile Map7D2 may enhance MT stabilization.

**Figure 4.**
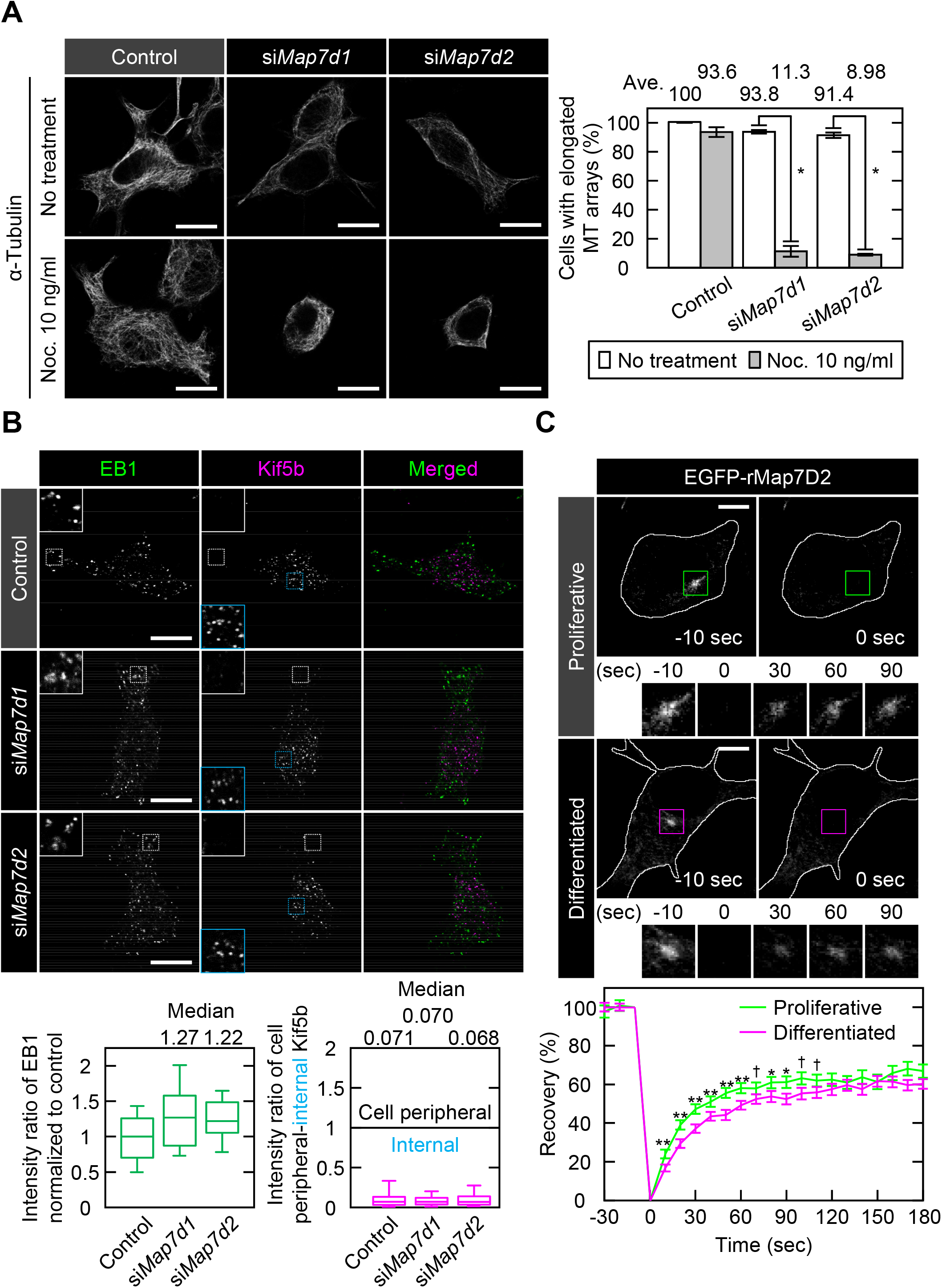
Map7D2 is required for MT stabilization within the cell. (**A**) Confirmation of MT stability by low-dose nocodazole treatment in the indicated N1-E115 cells. The cells were treated with or without a low concentration of nocodazole (10 ng/mL) for 1 h and were stained with an anti-α-tubulin antibody. Cells with the elongated MT arrays or the MT shrinkage were counted, and the rate of cells with the elongated MT arrays was calculated. Data are from three independent experiments and represent the average□±□SD. (**B**) Immunofluorescence staining for EB1 and Kif5b in N1-E115 cells treated with each siRNA (Top panels). The insets show enlarged images of regions indicated by a white box. Bottom left panel, the intensities of EB1 at the cell periphery in the indicated cells were measured via ROI analysis (each, n = 30 cells from three independent experiments). Data from *Map7d1* or *Map7d2* knock-down were shown by normalizing with the control value. Bottom right panel, the intensities of Kif5b at the cell periphery (white box) and the internal region (cyan box) were measured via ROI analysis, and the intensity ratios of cell peripheral-internal Kif5b were calculated (each, n = 30 cells from three independent experiments). Of note, a value of 1 means that Kif5b is distributed throughout the cell, and a value greater or less than 1 means that Kif5b is distributed at the cell periphery or in the internal region, respectively. (**C**) FRAP analysis of Map7D2 in proliferative or differentiated N1-E115 cells stably expressing EGFP-rMap7D2. Top and middle panels, recovery of fluorescence was recorded at 10-sec intervals for 3 min. Bottom panel, each recovery rate represents the median with SEM. Proliferative or differentiated state, n = 12 ROIs each from three independent experiments. Data information: In **(A)**, *, *P* < 0.0008 (the Student’s *t*-test). In **(C)**, **, *P* < 0.002, *, *P* < 0.01, †, *P* < 0.05 (the Student’s *t*-test). Scale bars, 10 μm in **(A-C)**.

The acetylation and detyrosination of α-tubulins are associated with stable MTs (Baas et al 2016, Janke & Montagnac 2017). Therefore, we examined the effects of *Map7d2* or *Map7d1* knock-down on the levels of acetylated and detyrosinated tubulins. Neither *Map7d2* knock-down nor knock-out affected the total levels of these modified tubulins (Fig. 5A and B). In contrast, *Map7d1* knock-down reduced the total level of acetylated but not detyrosinated tubulin (Fig. 5A), and double knock-down of *Map7d2* and *Map7d1* had the same effect (Fig. 5A). Consistent herewith, immunostaining revealed that *Map7d1* knock-down greatly decreased the intensity of acetylated tubulin around the centrosome in N1-E115 cells (Fig. 5C and D). *Map7d1* knock-down decreased the intensity of α-tubulin and increased that of Map7D2 (Fig. 5C and D), indicating that Map7D1 is required for the maintenance of acetylated, stable MTs. Under *Map7d2* knock-down, the intensity of α-tubulin and Map7D1 decreased without affecting that of acetylated tubulin (Fig. 5C and D). This decrease in the intensity of Map7D1 is presumably due to a reduction in the number of MT structures that can serve as scaffolds for Map7D1, because the total amount of Map7D1 in the cells is not affected by *Map7d2* knock-down or knock-out (Fig. 5A and B). Together with our biochemical data for Map7D2, these results suggest that Map7D2 facilitates MT stabilization via direct binding, in contrast to Map7D1.

**Figure 5.**
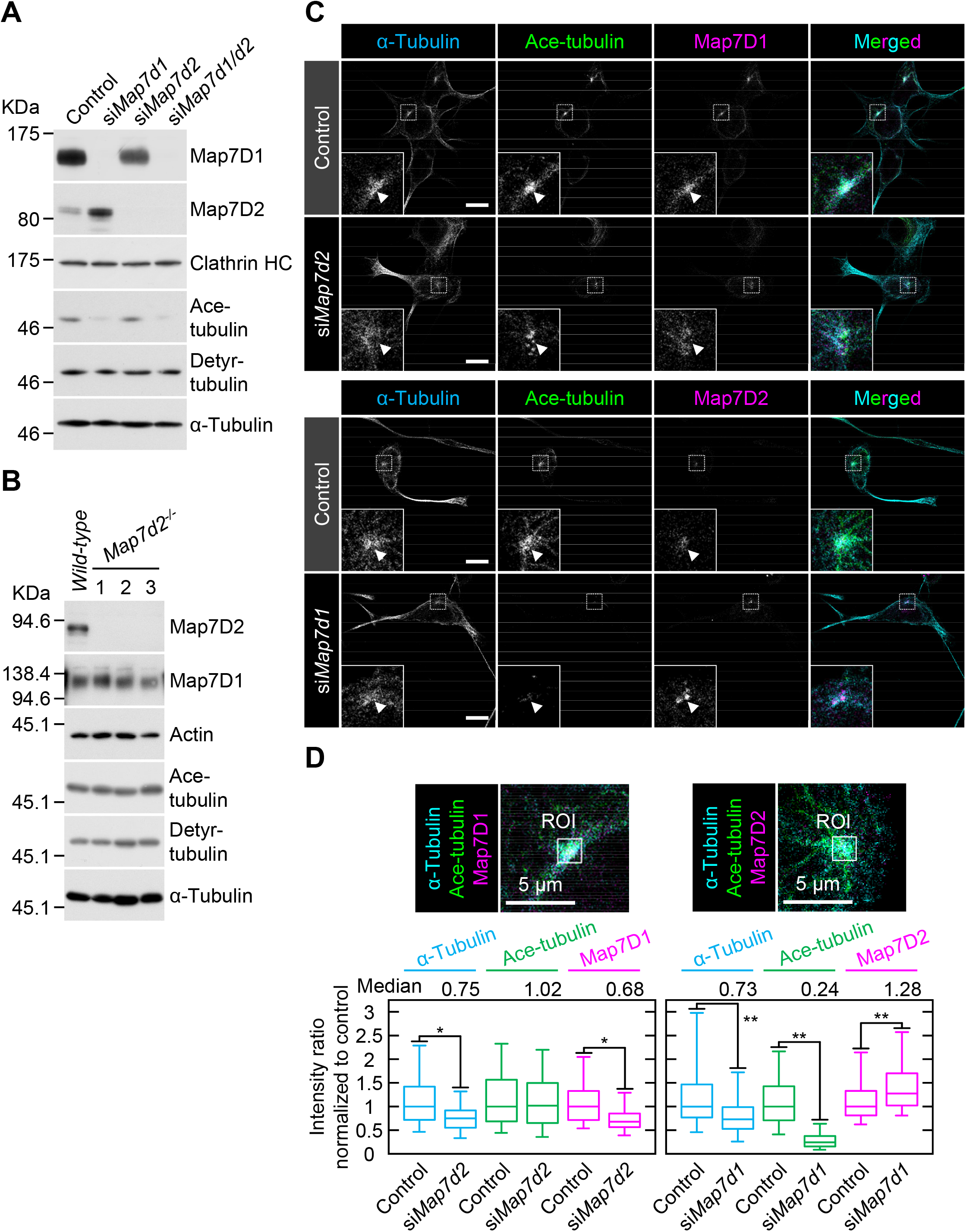
Map7D2 and Map7D1 facilitate MT stabilization through distinct mechanisms. (**A**) Immunoblot analysis for acetylated (Ace-) and detyrosinated (Detyr-) tubulin in N1-E115 cells treated with each siRNA. Lysates derived from the indicated cells were separated by SDS-PAGE and subjected to immunoblotting with anti-Map7D1, anti-Map7D2, anti-Ace-tubulin, or anti-Detyr-tubulin antibodies. The blot was reprobed for Clathrin heavy chain (HC) or α-tubulin as a loading control. (**B**) Immunoblot analysis for Ace- and Detyr-tubulins in *wild-type* and *Map7d2*^*-/-*^ N1-E115 cells. Three independent *Map7d2*^*-/-*^ clones were used in this study. Lysates derived from the indicated cells were separated by SDS-PAGE and were immunoblotted with anti-Map7D1, anti-Map7D2, anti-Ace-tubulin, or anti-Detyr-tubulin antibodies. The blot was reprobed for α-actin or α-tubulin as a loading control. (**C**) Immunofluorescence staining for α-tubulin, Ace-tubulin, and Map7D1 or Map7D2 in N1-E115 cells treated with each siRNA. For a comparison of signal intensities, images were captured under the same parameters. The insets show enlarged images of regions indicated by a white box. Of note, Ace-tubulin was present predominately around the centrosome in N1-E115 cells as indicated by arrowheads. (**D**) Quantification for immunofluorescence staining shown in (**C**). Left panels, the intensities of α-tubulin, Ace-tubulin, and Map7D1 around the centrosome in the indicated cells were measured via ROI analysis (control, n = 197 cells; si*Map7d2*, n = 192 cells from three independent experiments). Right panels, the intensities of α-tubulin, Ace-tubulin, and Map7D2 around the centrosome in the indicated cells were measured by ROI analysis (control, n = 193 cells; si*Map7d1*, n = 227 cells from three independent experiments). Data information: In **(D)**, the bars of box-and-whisker plots show the 5 and 95 percentiles. *, *P* < 1×10^−13^; **, *P* < 1×10^−8^ (the Student’s *t*-test). Scale bars, 10 μm in **(C)** and 5 μm in **(D)**.

Dysregulation of MT stabilization affects various biological functions. For instance, it can lead to increased cell motility and neurite outgrowth (Alesi et al 2016, Biernat et al 2002, Grenningloh et al 2004, Hubbert et al 2002). Therefore, we analyzed whether random cell migration or neurite outgrowth of N1-E115 cells would be affected by single or double knock-down of *Map7d2* and *Map7d1* and *Map7d2* knockout. As expected, each single knock-down and *Map7d2* knock-out enhanced not only the migration speed and distance during random cell migration (Fig. 6A-C), but also the rate of neurite outgrowth (Fig. 6D and Fig. S7C). Furthermore, double knock-down of *Map7d2* and *Map7d1* tended to result in increased cell motility and neurite outgrowth, compared with each single knock-down (Fig. 6B-D). Taken together, these results suggest that Map7D2 and Map7D1 facilitate MT stabilization through distinct mechanisms, thus controlling cell motility and neurite outgrowth.

**Figure 6.**
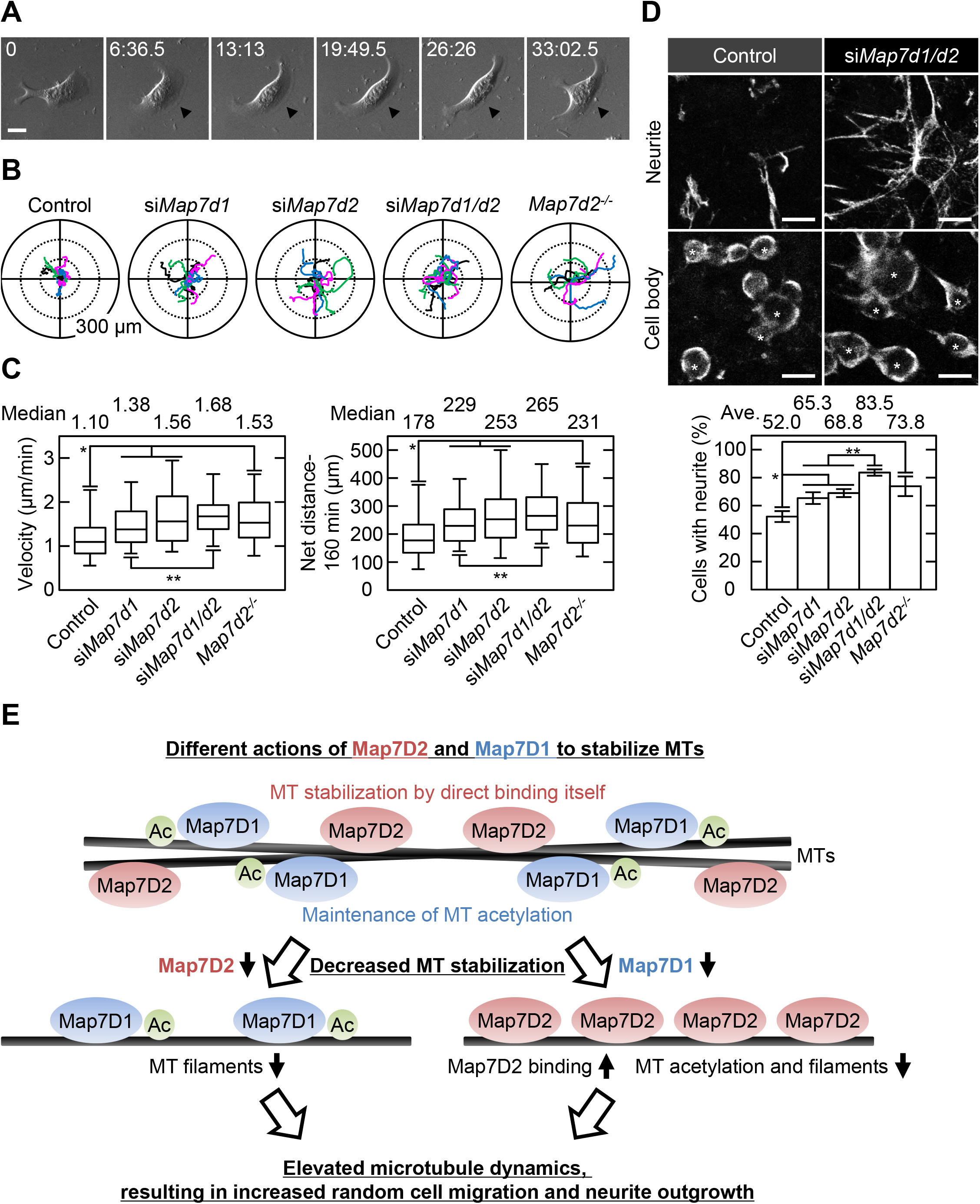
Map7D2 suppresses random cell migration and neurite outgrowth. (**A**) Bright-field images of migrating N1-E115 cells. Arrowheads show lamellipodia formed in the direction of migration. (**B**) Tracking analysis of random cell migration in the indicated cells. Each color represents the trajectory of 12 randomly selected cells. (**C**) Velocity and net distance measured in the indicated cells (control: n =114 cells; si*Map7d1*: n = 100 cells; si*Map7d2*: n = 71 cells; si*Map7d1/d2*: n = 107 cells; *Map7d2*^*-/-*^: n = 60 cells from three independent experiments). (**D**) Neurite outgrowth assay in the indicated cells. Neurites and cell bodies were visualized by α-tubulin staining (upper). The neurite outgrowth from each cell was distinguished by acquiring images with Z-sectioning. Data are from three or four independent experiments and represent the average□±□SD. (**E**) Proposed model for the distinct mechanisms of Map7D2 and Map7D1 for MT stabilization. See Discussion for further detail. Data information: In **(C)**, the bars of box-and-whisker plots show the 5 and 95 percentiles. *\*, P* < 1×10^−4^; **, *P* < 0.002 (the Student’s *t*-test). In **(D)**, *, *P* < 0.002; **, *P* < 0.0002 (the Student’s *t*-test). Scale bars, 20 μm in **(A, D)**.

## Discussion

In the present study, we provide the comprehensive analysis of Map7D2 biochemical properties (Fig. 2). The N-terminal and C-terminal regions of Map7D2 exhibited high homology to those of Map7 (Fig. S1). The N-terminal homologous region is basic and highly charged. Most MT-binding domains characterized thus far are confined to positively charged regions (Aizawa et al 1990, Lewis et al 1989, Noble et al 1989, Pierre et al 1992). Consistently, the MT-binding region of Map7 was shown to be located at the N-terminal positively charged region (Masson & Kreis 1993). Since we demonstrated that the N-terminal half of rMap7D2 directly bound to MTs (Fig. 2C), it is likely that Map7D2 also associates with MTs through the positively charged N-terminal region. On the other hand, the C-terminal region of Map7 includes the Kif5b binding domain (Metzger et al 2012), which mediates Kif5b-dependent transport by loading Kif5b onto MTs (Hooikaas et al 2019, Kikuchi et al 2018, Metzger et al 2012, Tymanskyj et al 2018). Consistent with the conservation of this region between Map7 and Map7D2, Map7D2 also has the ability to form a complex with Kif5b (Fig. S4) and contributes to Kinesin-1-mediated transport in the axons of hippocampal neurons (Pan et al 2019). Therefore, the biochemical properties of Map7D2 and Map7 are largely similar.

In contrast, the cellular functions of Map7D2 may differ from those of Map7. Our group and Hooikaas et al. have previously reported that Map7 and Map7D1 have functional overlaps in HeLa cells (Hooikaas et al 2019, Kikuchi et al 2018). For instance, both form a complex with Dishevelled, a mediator of Wnt5a signaling, while Map7D2 does not (Kikuchi et al 2018). In addition, Map7D2 exhibits distinct localization patterns in cultured hippocampal neurons, localizing to the proximal axon (Pan et al 2019). In the present study, we propose a molecular mechanism explaining how Map7D2 and Map7D1 regulate MT stabilization in N1-E115 cells (Fig. 6E). Map7D2 and Map7D1 both strongly localize to the centrosome and partially on MTs in proliferating as well as in differentiated N1-E115 cells (Fig. 3 and Fig. S5B-D). Further, the knock-down of either resulted in a comparable reduction of MT stabilization (Fig. 4A, Fig. 5C, and D), without affecting the amount of EB1-decorated MTs (Fig. 4B). Mechanistically, Map7D1 is required for the maintenance of MT acetylation, which is enriched in stable MTs, whereas Map7D2 is not (Fig. 4). While the above phenotypes exhibited by each knock-down are thought to enhance the rate of cell motility and neurite outgrowth, it is also possible that Map7D2 and Map7D1 control cell motility and neurite outgrowth through their function as complexes with Kif5b (Fig. S4) (Kikuchi et al 2018, Pan et al 2019). However, it has been reported that when Kinesin-1 function is impaired, both cell motility and neurite outgrowth are reduced (Agarwal et al 2019, Lu et al 2013). In addition, even after the knock-down of either *Map7d2* or *Map7d1*, the distribution of Kif5b foci was similar to that in control N1-E115 cells (Fig. 4B and Fig. S7A). Although it is possible that in *Map7d2* or *Map7d1* knock-down N1-E115 cells, the effects of reduced MT stabilization are offset by those of Kinesin-1 dysfunction, resulting in a mild increase in the rate of cell motility and neurite outgrowth, the phenotypes we observed are likely independent of the functions associated with Kif5b in N1-E115 cells. Alternatively, the fact that no stronger phenotype was observed may be because, besides Map7D2 and Map7D1, other molecules are involved in MT stabilization. Taking these findings and our biochemical data into consideration, we propose that, in contrast to Map7D1, Map7D2 facilitates stabilization by directly binding MTs, eventually to control cell motility and neurite outgrowth.

We also found that among the stable MT marker acetylated and detyrosinated tubulins, *Map7d1* knock-down reduced only acetylated tubulin (Fig. 5A, C, and D). These two modifications are mediated by different mechanisms and are sometimes not synchronous (Bance et al 2019, Song & Brady 2015, Sudo & Baas 2010, Yoshiyama et al 2003). For instance, Montagnac et al. have shown that defects in the α-tubulin acetyltransferase αTAT1-Clathrin-dependent endocytosis axis reduce only acetylated tubulin, resulting in a shift from directional to random cell migration (Montagnac et al 2013). In addition, acetylated tubulin is down-regulated by tubulin deacetylases such as HDAC6 and Sirt2 (Hubbert et al 2002, North et al 2003). Therefore, it is conceivable that Map7D1 may be involved in the pathway that controls the level of acetylated tubulin.

We also determined the tissue distribution of Map7D2, which has not been described to date (Fig. 1). Among the MAP7 family members, the tissue distribution of Map7 has been analyzed in detail so far (Fabre-Jonca et al 1998, Komada et al 2000). At the mRNA level, *Map7* is expressed in a variety of epithelial tissues, dorsal root ganglia, trigeminal ganglia, and primitive seminiferous tubules during embryonic development. We also reported that both Map7 and Map7D1 are expressed in the epithelia of the mouse fallopian tube at the protein level (Kikuchi et al 2018). Consistent with Map7 expression in primitive seminiferous tubules, *Map7* homozygous gene-trap mice exhibited defects in spermatogenesis (Komada et al 2000). Map7D2 was expressed predominantly in the glomerular layer of the olfactory bulb and Sertoli cells of the testis (Fig. 1C). The glomerular layer is known to be the region where axons accumulate and does not express Map2, a marker of neuronal cell bodies and dendrites (Fig. 1C). As Map7D2 localizes to the proximal axon in cultured hippocampal neurons (Pan et al 2019), Map7D2 may have similar localization and function in olfactory bulb neurons. The function of Map7D2 in Sertoli cells was not clarified in the present study. Therefore, whether Map7D2 is involved in mammalian neurogenesis and spermatogenesis represents a question for future research.

## Materials and methods

### Molecular cloning, expression, and purification of rMap7D2

Based on the information of DDBJ/EMBL/GenBank accession number XM_228973, oligonucleotide primers (5′-ATGTCGACATGGAGCGCAGCGGTGGGAACGGCG-3′ and 5-ATGTCGACTCAACAGAAGGTGTTCAGCGTAGTTTC-3′) were designed, and rat *Map7d2* (r*Map7d2*) cDNA was obtained by PCR using rat cDNA as a template. Expression vectors for r*Map7d2* were constructed in pCMV5-Myc (Nakanishi 1997), pQE9 (Qiagen), pGEX-5X-3 (Cytiva), pcDNA3.1/V5-His (Thermo Fisher Scientific), pCLXSN-GFP (Reiley et al 2005), and pEGFP-N3 (Clontech). GST-fused proteins were expressed in *Escherichia coli* and purified using glutathione-Sepharose beads (Cytiva), respectively. GST-rMap7D2 (full length) was further purified by gel filtration using a HiLoad 16/60 Superdex 200 column (Cytiva).

### Antibodies

A rabbit polyclonal anti-Map7D2 antibody was raised against GST-rMap7D2 (aa1-235). All the primary antibodies used are listed in Table S1. Secondary antibodies coupled to horseradish peroxidase (HRP) were purchased from Sigma-Aldrich. Alexa Fluor-conjugated secondary antibodies used for immunofluorescence experiments were purchased from Thermo Fisher Scientific.

### MT binding assay

The MT co-sedimentation assay was performed as previously described (Yamamoto 2002), with a slight modification. MTs were prepared by incubating tubulin in polymerization buffer (80 mM PIPES/NaOH [pH 6.8], 1 mM MgCl_2_, 1 mM EGTA, and 1 mM GTP) containing 10% glycerol for 20 min at 37 □. After incubation, Taxol was added at a final concentration of 15 μM. Various amounts of rMap7D2 were incubated with 0.4 mg/mL of MTs in polymerization buffer containing 15 μM taxol for 20 min at 37 □. After incubation with MTs, the mixture (200 μL) was placed over a 700-μL cushion of 50% sucrose in polymerization buffer containing 15 μM taxol. After the sample was centrifuged at 100,000 × g for 30 min at 37 □, the supernatant was removed from the cushion, and the original volume was restored with SDS sample buffer. Comparable amounts of the supernatant and pellet fractions were subjected to SDS-PAGE, followed by CBB protein staining. The amount of protein was estimated using a densitometer. ELISA for MT binding was performed in a 96-well microtiter plate as previously described (Pedrotti et al 1994). Briefly, wells were coated by incubating with 0.2 mg/mL of MTs in polymerization buffer containing 15 μM taxol for 2 h at 37 □ and then blocked via incubation with 5% glycine. Increasing amounts of rMap7D2 were added to each well and incubated for 20 min at 37 □. The plate was washed and further incubated with an anti-Map7D2 antibody. After washing, the plates were incubated with a secondary antibody conjugated to horseradish peroxidase. SuperSignal ELISA Pico (Pierce) was used as a chemiluminescent peroxidase substrate.

### MT polymerization assays

MT assembly was assayed by measuring turbidity at 350 nm using a spectrophotometer, as previously described (Gaskin et al 1974). Briefly, GST-rMap7D2 (0.14 mg/mL) was incubated with 2 mg/mL tubulin in polymerization buffer at 37 □. The sample was continuously monitored at 350 nm using a Hitachi U-2000 spectrophotometer. MT assembly was further assayed by fluorescence microscopy using rhodamine-labeled tubulin (Hyman 1991). Briefly, GST-rMap7D2 (0.07 mg/ml) was incubated at 37 □ for 20 min with 0.8 mg/mL tubulin (1 : 9 = rhodamine-labeled tubulin : unlabeled tubulin) in polymerization buffer. Incubation was stopped through the addition of 1% glutaraldehyde. The sample was spotted onto a glass slide and viewed under a fluorescence microscope.

### Northern blotting

An RNA blot membrane (Clontech) was hybridized with the ^32^P-labeled full coding sequence of rMap7D2, according to the manufacturer’s protocol.

### Cell culture and transfection

HeLa and N1-E115 cells were cultured at 37 □ in DMEM supplemented with 10% fetal bovine serum and penicillin-streptomycin. The methods employed for plasmid or siRNA transfection were previously described (Kikuchi et al 2010). Plasmid transfection of N1-E115 cells was performed using Lipofectamine LTX according to the manufacturer’s instructions. Differentiation of N1-E115 neuroblastoma cells was induced by decreasing the serum level to 0.5% fetal bovine serum and adding 1% dimethyl sulfoxide (DMSO) (hereafter, the above-described medium was referred to as differentiation medium). Stealth double-stranded RNA was purchased from Thermo Fisher Scientific (USA). All siRNAs used in this study are listed in Table S2. Three individual siRNAs for mouse *Map7d2* or *Map7d1* were designed based on the respective sequences. Double-stranded RNA targeting luciferase was used as a control. The cells were cultured for 72 h and subjected to various experiments. In Fig. 3D, Fig. 4A, B, Fig. 5A, C, D, and Fig. 6B-D, to exclude siRNA off-target effects, a mixture of three individual siRNAs for Map7D1 or Map7D2 was used. For the generation of N1-E115 cells stably expressing EGFP-rMap7D2, clones were selected by adding G418 at 24 h post-transfection. EGFP-rMap7D2 expression was confirmed by immunoblotting using antibodies against GFP and Map7D2. For Fig. 4A, cells were treated with 10 ng/mL of nocodazole (Sigma-Aldrich) for 1 h.

### Generation of *Map7d2* knock-out N1-E115 cell lines by CRISPR-Cas9

Two sgRNA sequences were designed using the CHOPCHOP CRISPR/Cas9 gRNA finder tool (http://chopchop.cbu.uib.no/). The short double-stranded DNA for each sgRNA (5-CACCGTGAAGAGAGCACATGTGCC-3 and 5-AAACGGCACATGTGCTCTCTTCAC-3, or 5-CACCGCAGGATCACCAGGGCCTGG-3 and 5-AAACCCAGGCCCTGGTGATCCTGC-3′) were inserted into the *Bbs*I site of pX330 (Cong et al 2013). To construct the *Map7d2* knock-out vector, the 5′ and 3′ arms of each gene were amplified by PCR using N1-E115 genomic DNA and cloned into the pCR4 Blunt-TOPO vector (Thermo Fisher Scientific). The puromycin resistance marker was inserted between the 5′ and 3′ arms (Fig. S6A). N1-E115 cells were transfected with 1 μg of each of the two pX330-sgRNA plasmids and the knock-out vector using Lipofectamine LTX (Thermo Fisher Scientific). Knock-out clones were selected by adding puromycin (Sigma-Aldrich) at 24 h post-transfection. Successful knock-out was confirmed by immunoblotting using an anti-Map7D2 antibody and genomic PCR.

### Animals

Mice (C57BL6/N; Japan SLC, Japan) were used in this study. Animal care and experiments were conducted in accordance with the guidelines for the care and use of laboratory animals of the Center for Animal Resources and Development, Kumamoto University. All experiments were approved by the experimental animal ethics committee of Kumamoto University (A2019-127 and A2021-018). Mice were kept in a light- and temperature-controlled room with a 12-h light/dark cycle at 22 ± 1 □.

### Quantitative real-time PCR

Each RNA sample was subjected to reverse transcription using murine leukemia virus reverse transcriptase (Thermo Fisher Scientific), and the generated cDNA was used as a template for qRT-PCR. Each reaction mixture was prepared using the KAPA SYBR Fast qPCR kit (Kapa Biosystems), and the PCR reaction was performed on ViiA7 (Thermo Fisher Scientific). The primers used for RT-qPCR are listed in Table S3.

### Immunoblotting and immunoprecipitation

For immunoblotting, cells were washed once with PBS and lysed with Laemmli’s sample buffer. After boiling, the lysates were separated by SDS–PAGE, transferred to PVDF membranes (Millipore), and immunoblotted with antibodies. For immunoprecipitation analysis, the HeLa cells were washed once with PBS at 24 h post-transfection and lysed with 1× NP40 buffer [20 mM Tris-HCl (pH 8.0), 10% glycerol, 137 mM NaCl, 1% NP40] supplemented with protease inhibitors and phosphatase inhibitors for 20 min on ice. The supernatant was collected after centrifugation and incubated with the appropriate antibodies. After incubation, 15 μL of protein A or G Sepharose beads was added, and the mixtures were rotated for 1 h at 4 □. The beads were washed once with 1×NP40 buffer, twice with LiCl buffer [0.1 M Tris-HCl (pH 7.5), 0.5 M LiCl], once with 10 mM Tris-HCl (pH 7.5), and were finally resuspended in Laemmli’s sample buffer.

### Immunofluorescence and imaging analyses

For immunofluorescence staining, cells were grown on coverslips and fixed in 100% methanol at -20 □ for 5 min. After blocking with 1% BSA in PBS for 1 h at room temperature, the samples were incubated with primary antibodies overnight at 4 □, followed by incubation with Alexa Fluor-conjugated secondary antibodies (Thermo Fisher Scientific) for 1 h. For immunofluorescence tissue staining, tissues were fixed in 4% paraformaldehyde in PBS at 4 □ overnight, and then immersed sequentially in 10, 20, and 30 % sucrose in PBS at 4 □. After sucrose equilibration, tissues were immersed in OCT (Sakura Finetechnical) at room temperature for 5 min, followed by embedding in OCT and freezing in liquid nitrogen. Sections (10 μm) were stored at −80 □. The sections were washed once with PBS for 10 min and twice with 0.1 % Triton X-100 in PBS for 10 min. After blocking with Blocking One (Nacalai) for 1 h at room temperature, the samples were incubated with primary antibodies overnight at 4 □, followed by incubation with Alexa Fluor-conjugated secondary antibodies (Thermo Fisher Scientific) for 1 h. Nuclei were stained with DAPI for 30 min at room temperature. The samples were viewed under a fluorescence microscope (Olympus, BX51) or a confocal microscope (Olympus, FV1000 or Leica, TCS SP8). For Fig. 4C, FRAP analysis in N1-E115 cells stably expressing EGFP-rMAp7D2 were performed with an FV1200 equipped with GaAsP detectors and an incubator (Olympus). Images were processed and analyzed using Fiji software (National Institutes of Health).

### Random cell migration assay and neurite outgrowth assay

For the random cell migration assay, cells were seeded onto a laminin-coated (10 μg/mL) glass-bottom dish and recorded under an inverted microscope system equipped with an incubator (Olympus, LCV110). For the neurite outgrowth assay (Fig. S7C), the underside of 3 μm pore transwell membranes (Corning) was coated with 500 μL of 10 μg/mL laminin in PBS into a well of a 24-well plate. After coating, the membranes were removed from the laminin and placed into the well of a 24-well dish containing 500 μL differentiation media. One hundred microliters of cell suspension (containing 1–2 × 10^5^ cells) was added to the insert chamber on top of the membrane. The cells were allowed to extend neurites through the membrane pores to the lower chamber (underside of the membrane) for 6 h at 37 □. The cells were then fixed and stained with an anti-α-tubulin antibody. Images were processed and analyzed using Fiji software (National Institutes of Health).

### Statistics

The experiments were performed at least three times (biological replicates), and the results are expressed as the average ± S.D. or the median, first and third quartiles, and 5-95 % confidence intervals for the box-and-whisker plot. Differences between data values, except for Fig. 2D, were tested for statistical significance using the Student’s *t*-test. Statistical significance was set at *P* <0.05. In Fig. 2D, differences between data values were tested for statistical significance using the F-test.

### Other Procedures

Tubulin was prepared from fresh porcine brains by three cycles of polymerization and depolymerization, followed by DEAE-Sephadex column chromatography (R.C.Jr. & Lee 1982, Shelanski et al 1973).

## Data Availability

Source data for each figure are available in PDF or Excel format. In this study, we do not deposit any data in public databases.

## Acknowledgements

We thank past and present members of our laboratory for helpful discussions. This work was partly carried out at the Institute of Molecular Embryology and Genetics, the Gene Technology Centre, the Center for Animal resources and Development, and Research Facilities of the School of Medicine, Kumamoto University.

## Author Contributions

K.K. designed the research and carried out most of the experiments, except for Fig. 1A, B, and 2. Y.S., A.U., H.Y., and H.N. carried out Fig. 1A, B and 2, and raised the rabbit polyclonal anti-Map7D2 antibody. K.I. provided technical assistance for the immunofluorescence analysis using the mouse testis. K.S. provided technical assistance for the immunofluorescence analysis using the mouse brain. Y.S., T.S., S.H., and H.N. provided reagents, materials, and analysis tools. K.K. wrote the paper, and Y.S., and H.N. edited the paper. All of the authors discussed the results and commented on the manuscript.

## Figure legends

**Figure S1 - Sequence alignment among human MAP7 family proteins**.

*(A)* Schematic structures of human MAP7 family proteins, Map7, Map7D1, Map7D2, and Map7D3. (**B**) Sequence alignment of *conserved N-terminal regions in each* human MAP7 family protein. (**C**) Sequence alignment of *conserved C-terminal regions in each* human MAP7 family protein.

**Figure S2 - Confirmation for the specificity of anti-Map7D2 antibody**.

(**A**) Confirmation for the specificity of anti-Map7D2 antibody using V5His_6_-tagged human Map7 (hMap7) or rat Map7D2 (rMap7D2). An empty vector, hMap7-V5His_6_, or rMap7D2-V5His_6_ was transfected into HeLa cells, and cell lysates were subjected to SDS-PAGE, followed by immunoblotting with the anti-Map7D2 or anti-V5 antibody. (**B**) Endogenous expression of the MAP7 family genes in N1-E115 cells. Total RNA was extracted from N1-E115 cells and was subjected to reverse transcription. qPCR was performed with gene-specific primes using the cDNA as a template. Relative target gene mRNA expression was normalized to *Gapdh* expression. Data are from three independent experiments, and represent the average□±□SD. (**C**) Knock-down of endogenous *Map7d2* or *Map7d1* in N1-E115 cells. Cells were treated with each siRNA, and cell lysates were subjected to SDS-PAGE, followed by immunoblotting with the anti-Map7D2, anti-Map7, or anti-Clathrin HC antibody.

**Figure S3 - Database-based analyses for the expression distribution of *Map7d2* in the mouse brain**.

The *Map7d2* expression distribution in the mouse brain was analyzed using the dataset of RNA-Seq CAGE (left), RNA-Seq (middle), and SILAC (right) datasets in the Expression Atlas (https://www.ebi.ac.uk/gxa/home/RNA-seq).

**Figure S4 - Map7 or Map7D2 forms a complex with Kif5b, a member of Kinesin-1**.

Lysates from HeLa cells co-expressing hMap7-V5His_6_ (1), V5His_6_-tagged chimeric protein of rMap7D2 N-terminus and hMap7 C-terminus (2), or rMap7D2-V5His_6_ (3) with hKif5b-EGFP were immunoprecipitated with an anti-V5 antibody, and the immunoprecipitates were probed with anti-GFP and anti-V5 antibodies. E represents an empty vector control. Map7 has a region on the C terminus that is required for complex formation with Kif5b, a member of Kinesin-1.

**Figure S5 - Subcellular localization of Map7D1 in proliferative and differentiated N1-E115 cells**.

(**A**) Neuronal differentiation of N1-E115 cells was induced by decreasing the concentration of fetal bovine serum in the medium to 0.5% fetal bovine serum and adding 1% dimethylsulfoxide. Arrowheads indicate elongated neurites. After 6 h of induction, elongated neurites were observed. (**B-D**) Localization of Map7D1 in interphase (**B**), mitosis (**C**), and differentiation state

(**D**) of N1-E115 cells. Cells were double-stained with anti-Map7D1 and anti-α-tubulin antibodies. In **B**, the insets show enlarged images of regions indicated by a white box. In **C**, the inset shows metaphase cells. In **D**, images of differentiated cells were captured by z-sectioning, because the focal planes of the centrosome and neurites are different. Each inset shows an enlarged image of the region indicated with a white box at each focal plane. Arrow heads show centrosomal localization of Map7D1.

Data information: Scale bars, 50 μm in (**A**) and 10 μm in (**B**-**D**).

**Figure S6 - Generation of *Map7d2* knock-out N1-E115 cells**.

(**A**) Schematic representation of *Map7d2* knock-out (*Map7d2*^*-/-*^) N1-E115 cell generation using the CRISPR-Cas9 technique. (**B**) *Map7d2* knock-out was confirmed by immunoblotting. Three independent clones of *Map7d2*^*-/-*^ cells were used in this study. Lysates derived from the indicated cells were probed with anti-Map7D2 antibody. In addition, the effect of *Map7d2* knock-out on Map7D1 expression was analyzed using the anti-Map7D1 antibody. The blot was reprobed for α-tubulin as a loading control. The expression levels of the indicated proteins in *Map7d2*^*-/-*^ cells were carefully compared with dilution series of wild-type lysates.

**Figure S7 - Map7D2 stabilizes MTs within the cell**.

(**A**) Images of EB1 and Kif5b at the protrusion in N1-E115 cells treated with each siRNA, related to Fig. 4A. (**B**) Immobile fraction of Map7D2 at the proliferative or differentiated state of N1-E115 cells, related to Fig. 4C. From Fig. 4C, each immobile fraction of Map7D2 was calculated by subtracting each recovery rate from 100%. (**C**) Schematic representation of neurite outgrowth assay, related to Fig. 6D.

Data information: Scale bars, 10 μm in (**A**).

