## Supplementary material for "Map7D2 and Map7D1 facilitate microtubule stabilization through distinct mechanisms in neuronal cells": Figure S1

### Kikuchi_Figure S1

**A**

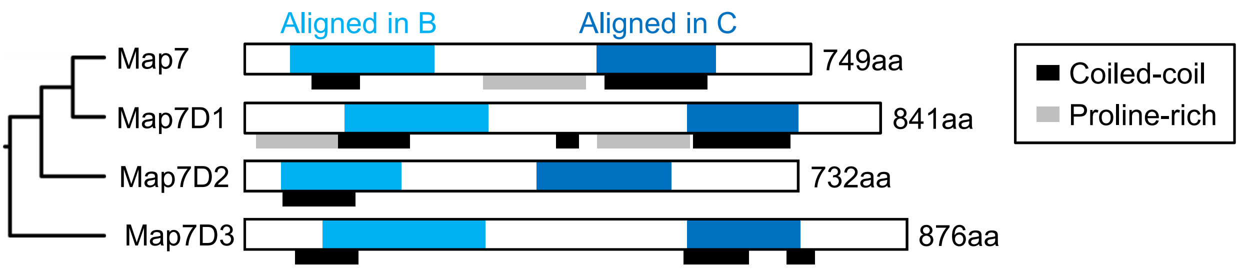


**B**

```
                 10        20        30        40        50        60
Map7    VLRVDDRQRLARERREEREKQLAAREIVWLEREERARQHYEKHLE---ERKKRLEEQRQK
Map7D1  VKKAGERHKLAKERREERAKYLAAKKAVWLEKEEKAKALREKQLQ---ERRRRLEEQRLK
Map7D2  FLKSDERQRLAKERREEREKCLAAREQQILEKQKRARLQYEKQME---ERWRKLEEQRQR
Map7D3  EKQMEERQRKLKERKEKEEQRRIAAEEKRHQKDEAQKEKFTAILYRTLERRRLADDYQQK

                 70        80        90       100       110       120
Map7    ---------------EERRRAAVEEKRRQ---RLEEDKERHEAVVRRTMERSQKPKQKHN
Map7D1  ---------------AEQRRAALEERQRQ---KLEKNKERYEAAIQRSVKKTWAEIRQQ-
Map7D2  ---------------EDQKRAAVEEKRKQ---KLREEEERLEAMMRRSLERTQQLELKK-
Map7D3  RWSWGGSAMANSESKTANKRSASTEKLEQGTSALIRQMPLSSAGLQNSVAKRKTDKERSS

                130       140       150       160       170       180
Map7    RWSWG-GSLHGSPSIHSADPDRRSVSTMNLSKYVDPVISKRLSSSSATLLNSPDRARRL-
Map7D1  RWSWA-GALHHSSPGHKTSGSRCSVSAVNLPKHVDSIINKRLSKSSATLWNSPSRNRSL-
Map7D2  KYSWG-APLAIGPGGHDAC-DKLSTSTMSLPKPTEPPMNKRLSSSTVAISYSPDRVFHVC
Map7D3  SLNRRDSNLHSSTDKEQAERKPRVTGVTNYVMQYVTVPLRKCTSDELRAVMFPMSTMKIP

                190       200       210
Map7    -QLSPWESSVVNRLLTPTHSFLARSKSTAALSGEA
Map7D1  -QLSAWESSIVDRLMTPTLSFLARSRSAVTLPRNG
Map7D2  P-----------RL---------------------
Map7D3  PQTKVEESPLEKVETPPKASVDAPPQVNVEVFCNT
```

**C**

```
                 10        20        30        40        50        60
Map7    ASVKTSAGTTDPEEATRLLAEKRRLAREQREKEERERREQEELERQKREELAQRVAEERT
Map7D1  TPSKPMAGTTDREEATRLLAEKRRQAREQREREEQERRLQAERDKRMREEQLAREAEARA
Map7D2  ALGKPTAGTTDAGEAAKILAEKRRQARLQKEQEEQERLEKEEQDRLEREELKRKAEEERL
Map7D3  SGNKSTAGIMNAEAATKILTELRRLAREQREKEEEERQREEMQQRVIKKSKDMAKEAVGG

                 70        80        90       100       110       120
Map7    TRREEESRRLEAEQAREKEEQLQRQ-------------------AEERALREREEAERAQ
Map7D1  EREAEARRREEQE-------------------------------AREKAQAEQEEQERLQ
Map7D2  RLEEEARKQEEERKRQEEEKKKQEGEEKRKAGEEAKRKAEEELLLKEKQEQEKQEKAMIE
Map7D3  QAEDHLKLKDGQQQNE----------------------------TKKKKGWLDQEDQEAP

                130       140       150       160       170
Map7    RQKE-EEARVREEAERVRQEREKHFQREEQERLERKKRLEEIMKRTRRTEATDKKTSD
Map7D1  KQKEEAEARSREEAERQRLEREKHFQQQEQERQERRKRLEEIMKRTRKSEVSETKQKQ
Map7D2  KQKEAAETKAREVAEQMRLEREQIMLQIEQERLERKKRIDEIMKRTRKSDVSPQVKKE
Map7D3  LQKGDAKIKAQEEADKRKKEHERIMLQNLQERLERKKRIEEIMKRTRKTDVNASKVTE
```
