## Supplementary figures and images for "Map7D2 and Map7D1 facilitate microtubule stabilization through distinct mechanisms in neuronal cells"

### Figure S2

Kikuchi_Figure S2

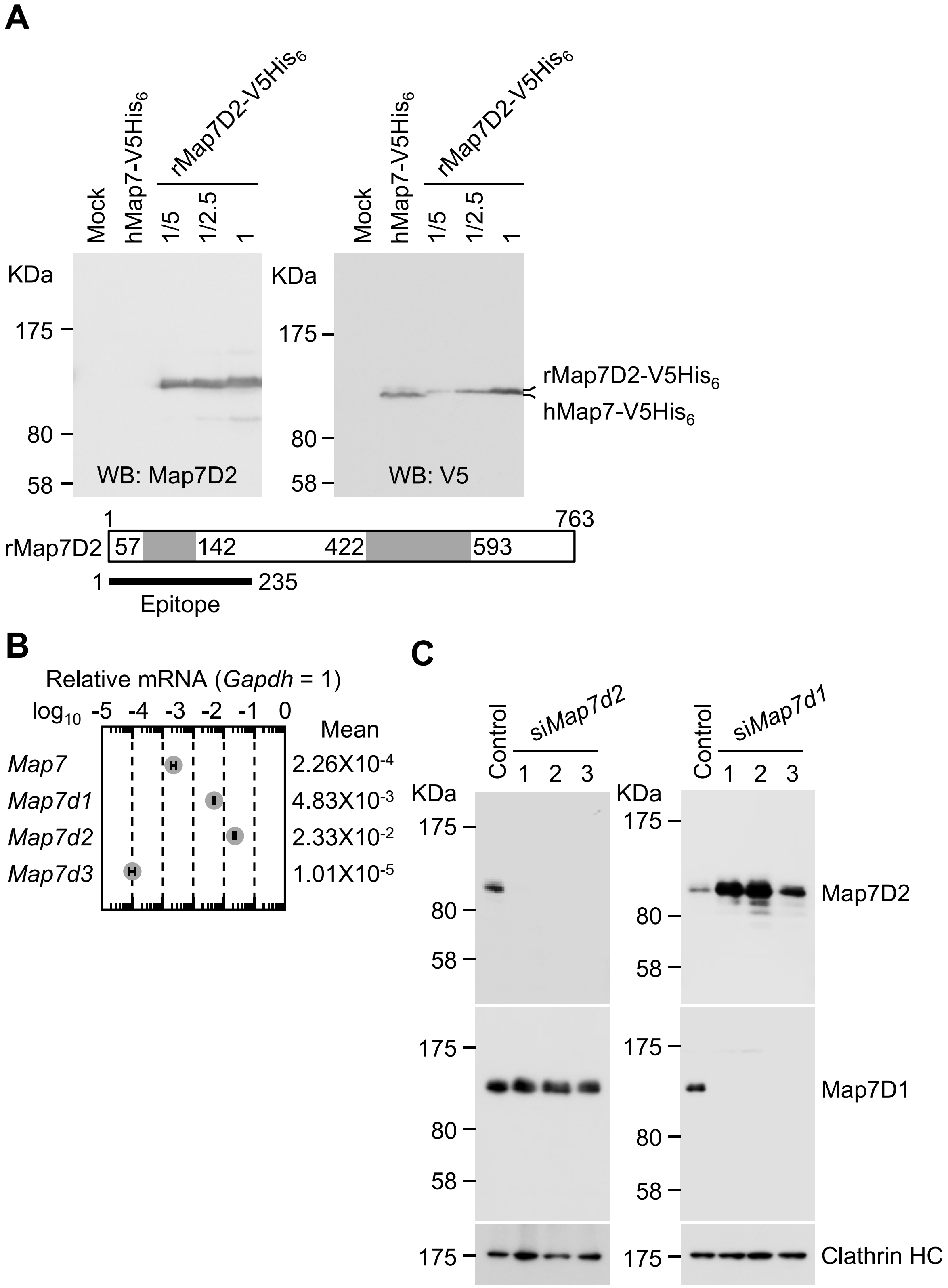

### Figure S3

Kikuchi_Figure S3

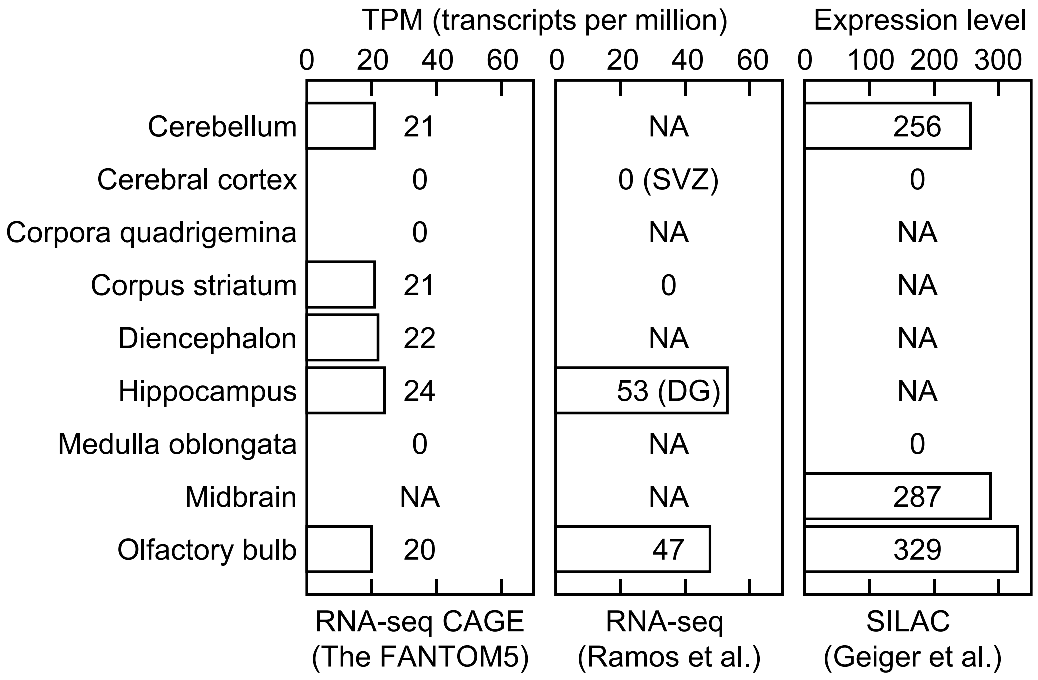

### Figure S4

#### Kikuchi_Figure S4

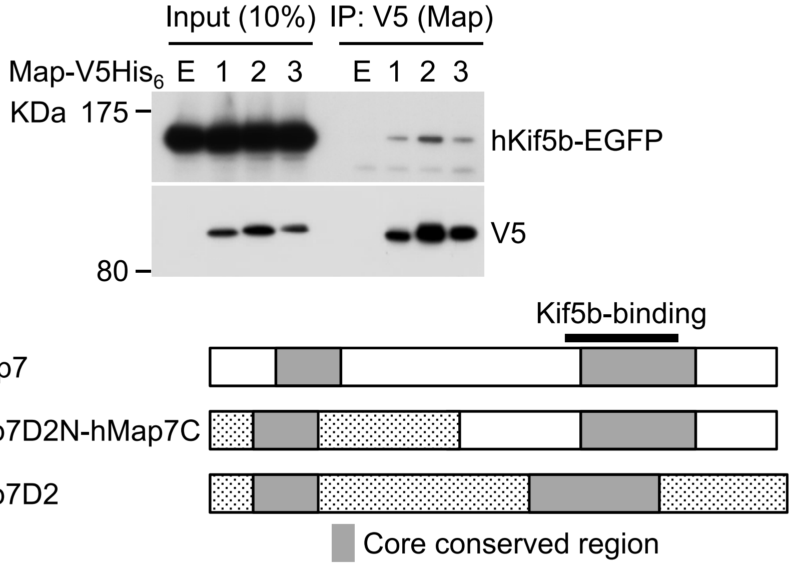

### Figure S5

Kikuchi_Figure S5

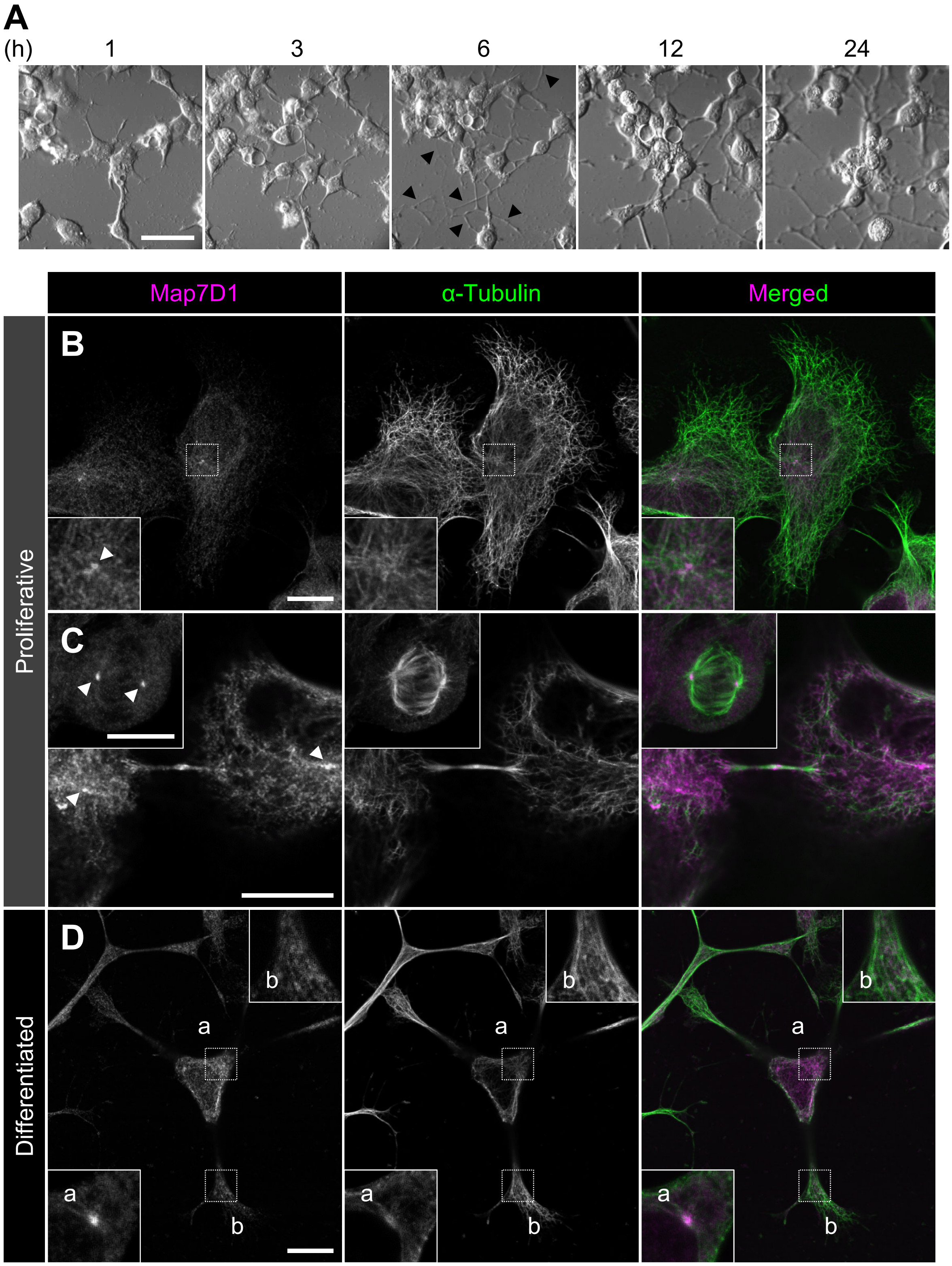

### Figure S6

Kikuchi_Figure S6

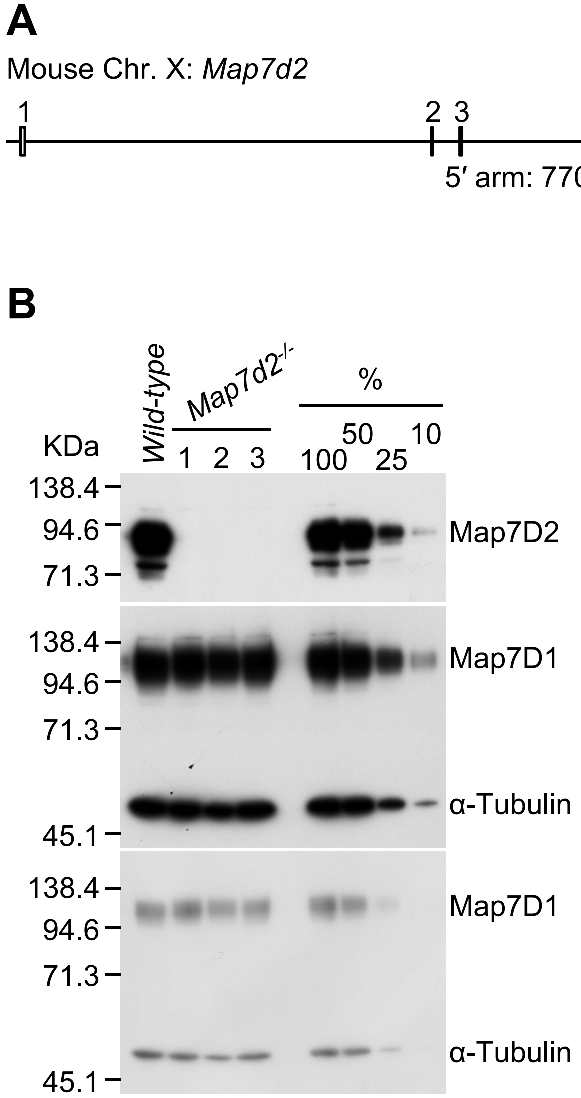

### Figure S7

Kikuchi_Figure S7

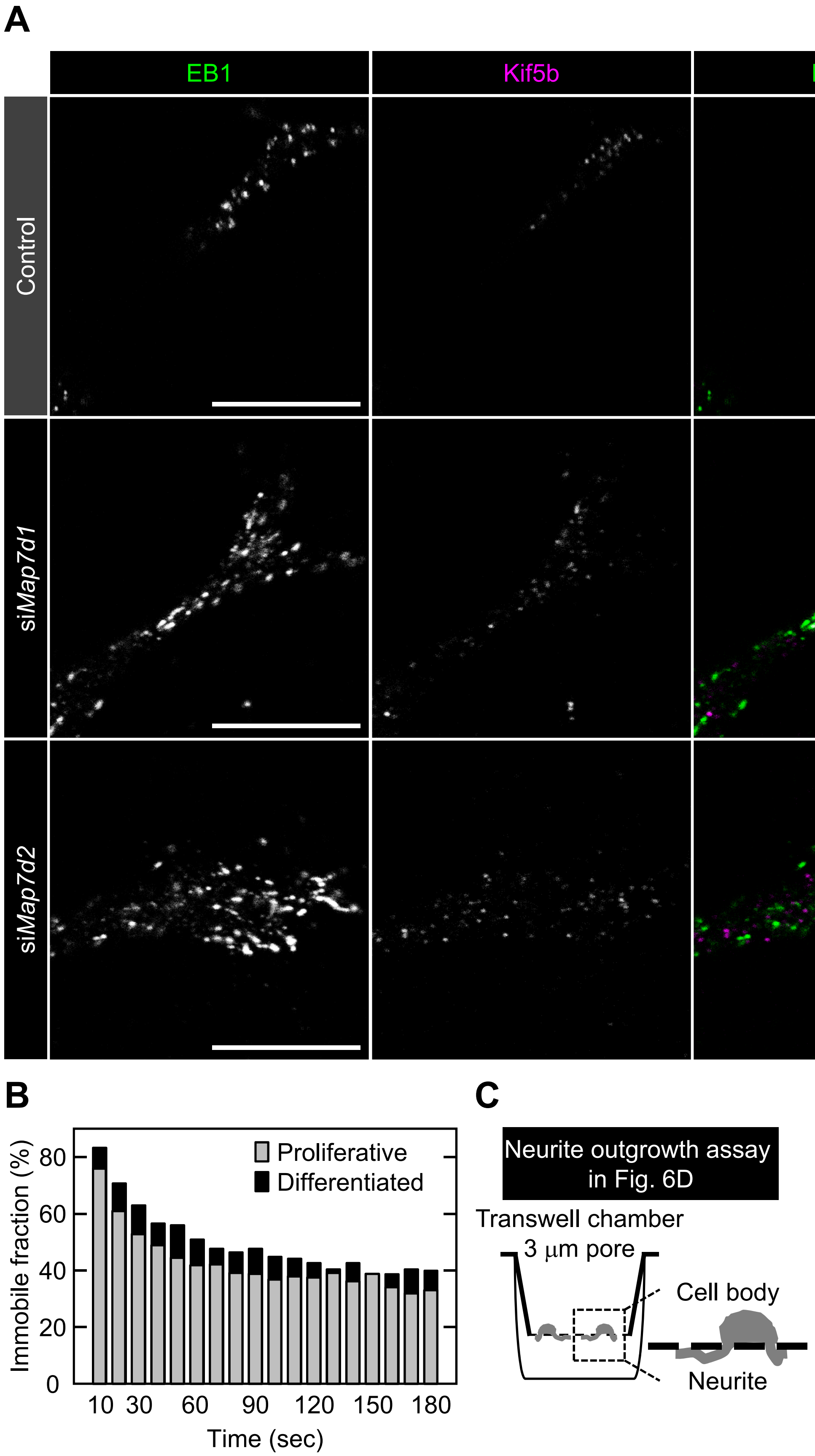
