## Supplemental information for "Map7D2 and Map7D1 facilitate microtubule stabilization through distinct mechanisms in neuronal cells"

**Table S1 - Primary antibodies used in this study.**

| Company | Name, catalog number | Used for (dilutions) |
| --- | --- | --- |
| BD Biosciences | Mouse anti-Clathrin heavy chain, 610500 | IB (1:5000) |
|  | Mouse anti-EB1(5/EB1), 610534, | IF (1:500) |
| GeneTex | Rabbit anti-Kif5b, GTX104874 | IF (1:500) |
| Millipore | Rabbit anti-Detyrosinated tubulin, AB3201 | IB (1:3000) |
| MP Biomedicals | Mouse anti-Actin (C4), 0869100-CF | IB (1:10000) |
| Nacalai | Mouse anti-V5, 04434-36 | IB (1:3000) |
|  | Rat anti-GFP, 04404-84 | IB (1:3000) |
| Sigma-Aldrich | Mouse anti-Acetylated tubulin, T7451 | IB (1:5000), IF (1:250) |
|  | Mouse anti-α-tubulin (DM1A), T6199 | IB (1:10000), IF (1:500) |
|  | Mouse anti-γ-tubulin (GTU-88), T6557 | IB (1:5000) |
|  | Mouse anti-Map2, | IF (1:250) |
|  | Mouse anti-Tubb3 | IF (1:250) |
| Made in-house | Rabbit anti-Map7D1 (Kikuchi et al 2018) | IB (1:10000), IF (1:500) |
|  | Rabbit anti-Map7D2 | IB (1:10000), IF (1:500) |
|  | Mouse anti-Myc monoclonal (9E10) | IF (1:250) |

IB, Immunoblotting; IF, Immunofluorescence

**Table S2 - siRNAs used in this study.**

| Name | Sequences |
| --- | --- |
| si*Luc* (+) | CGUACGCGGAAUACUUCGAAAUGUC |
| si*Luc* (-) | GACAUUUCGAAGUAUUCCGCGUACG |
| si*Map7d1*-1_380 (+) | UUUACAUCUUGCUUGGCAGGAGGGC |
| si*Map7d1*-1_380 (-) | GCCCUCCUGCCAAGCAAGAUGUAAA |
| si*Map7d1*-2_2144 (+) | UUCAUUAUCUCCUCCAGCCGCUUUC |
| si*Map7d1*-2_2144 (-) | GAAAGCGGCUGGAGGAGAUAAUGAA |
| si*Map7d1*-3_2157 (+) | UUUCCGAGUCCUCUUCAUUAUCUCC |
| si*Map7d1*-3_2157 (-) | GGAGAUAAUGAAGAGGACUCGGAAA |
| si*Map7d2*-1_1898 (+) | AACACAUUGAUUUCGAUCUUGUUGG |
| si*Map7d2*-1_1898 (-) | CCAACAAGAUCGAAAUCAAUGUGUU |
| si*Map7d2*-2_2062 (+) | UAGACUGAACGUCUUCAGUUGAGUC |
| si*Map7d2*-2_2062 (-) | GACUCAACUGAAGACGUUCAGUCUA |
| si*Map7d2*-3_2320 (+) | AACAGAAGGUAUUCAGGGUAGUUUC |
| si*Map7d2*-3_2320 (-) | GAAACUACCCUGAAUACCUUCUGUU |

**Table S3 - Primers for RT-qPCR in N1E115 cells.**

| Name | Sequences |
| --- | --- |
| m*Gapdh*_264 (+) | CGGTGCTGAGTATGTCG |
| m*Gapdh*_427 (-) | TGAGTGAGTTGTCATATTTCTCG |
| m*Map7*_983 (+) | TCAAAGCGAGGTCACCG |
| m*Map7*_1136 (-) | CGGATGTTGCCAGGAGA |
| m*Map7d1*_1304 (+) | GCGAACGGAACCTCAAGA |
| m*Map7d1*_1476 (-) | TGGAGGAAGGGCATGTC |
| m*Map7d2*_1098 (+) | TGCAAACGAAGAAACACCAA |
| m*Map7d2*_1257 (-) | CTTATCTGCTACATACTTCTCGG |
| m*Map7d3*_1074 (+) | ATCATTCTCCTTTGGGAGTGTA |
| m*Map7d3*_1252 (-) | CACTTGCTTCAAGGCGT |
